# Detecting latent gene-environment interaction when analyzing binary traits

**DOI:** 10.1101/2024.07.10.602954

**Authors:** Ziang Zhang, Jerald F. Lawless, Andrew D. Paterson, Lei Sun

## Abstract

In genome-wide association studies (GWAS), it is desirable to test for interactions (*GxE*) between single-nucleotide polymorphisms (SNPs,*G*’s) and environmental variables (*E*’s). However, directly accounting for interaction is often infeasible, because *E* is latent. For quantitative traits (*Y*) that are approximately normally distributed, it has been shown that indirect testing on *GxE* can be done by testing for heteroskedasticity of *Y* between genotypes. However, when traits are binary, the existing methodology based on testing the heteroskedasticity of the trait across genotypes cannot be generalized. In this paper, we propose an approach to indirectly test *GxE* for binary traits based on the non-additive effect *G*, and subsequently propose a joint test that accounts for the main and interaction effects of each SNP during GWAS. We illustrate the statistical features including type-I-error control and power of the proposed method through extensive numerical studies. Applying our method to the UK Biobank dataset, we showcase the practical utility of the proposed method, revealing SNPs and genes with strong potential for latent interaction effects.

## 1. Introduction

It is well known that the interaction (denoted as *GxE*) between single-nucleotide polymorphisms (SNPs; *G*’s) and environmental factors (*E*’s), or between SNPs (denoted as *GxG*), play an important role in shaping human complex traits (*Y*’s) (Manolio et al., 2009). A classic *GxE* example is the interaction effect between genetic variants in *PAH* and diet on the risk of phenylketonuria and its subsequent intellectual disability (Johns Hopkins University, 2024). Examples of *GxG* have also been reported Singhal et al. (2023).

However, a direct, exhaustive *G*x*G* search may be undesirable in the genome-wide association study (GWAS) setting because of the large-scale (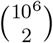 or more) multiple hypothesis testing. A direct *G*xE analysis, on the other hand, maybe infeasible in practice if the interacting *E* is latent or missing. For these reasons, it is often useful to conduct latent interaction analyses in GWAS; to simplify the notations, we use *GxE* hereinafter for both *GxE* and *GxG* scenarios.

For a quantitative trait *Y* that is approximately normally distributed, it has been shown that a latent *E* (or an un-modeled genetic variant) that interacts with a bi-allelic SNP *G* will produce heteroskedasticity in *Y* across the three genotypes of the SNP (Paré et al., 2010). Consequently, SNPs for which traits have shown significant heteroskedasticity (variance-quantitative trait loci, vQTLs) can be used to screen for potential *GxE*, and multiple vQTL methods have been developed (Young et al., 2018; Wang et al., 2019; Marderstein et al., 2021; Soave et al., 2015; Soave and Sun, 2017; Miao et al., 2022). The vQTL latent interaction approach has identified promising SNPs for follow-up interaction analysis. For example, rs12753193 near *LEPR* was first identified through vQTL analysis with evidence of interaction effect with BMI on C-reactive protein levels (Paré et al., 2010).

The lack of a corresponding latent *GxE* method for binary traits causes us to potentially miss novel findings. However, the variance technique for a quantitative trait cannot be used for a binary *Y*, as the variance of a binary trait is determined by its mean. Similarly, an over-dispersion parameter cannot be identified when *Y* is binary (Hinde and Demétrio, 1998). Thus, how to indirectly detect latent *GxE* in the binary setting remains an open problem.

In this paper, we first show that for binary traits commonly analyzed through the logistic and probit regression models, the latent *GxE* can be indirectly tested through the non-additive effect of *G*. Analogous to the joint location-scale test for a quantitative trait that integrates the vQTL information with the traditional mean-based GWAS (Soave et al., 2015), we then show how the joint test for a binary trait is related to the so-called genotypic test. Finally, we demonstrate the validity, power and practical applicability of the proposed method through extensive numerical studies and real data application.

## 2. Preliminary

### Indirect Test of Latent *GxE* for Quantitative Traits

Let *Y* be the trait of interest and *G* the genotypes of the SNP of interest, with the major and minor alleles coded as *a* and *A*, respectively, and the corresponding allele frequencies of *q* = 1 *− p* and *p* (*≤* 0.5), respectively; *p* is the minor allele frequency (MAF). Furthermore, let *G*_*A*_ denote the count of minor alleles *A* at a SNP, then *G*_*A*_ = 0, 1 and 2 corresponds to *G* = *aa, Aa* and *AA*, respectively, also termed additive coding.

If the environmental variable *E* is observed and hypothesized to interact with *G*, then the following linear regression would typically be used:

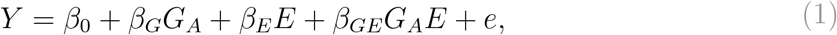

where 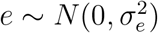 is independent of *G* and *E*, and *G* is typically assumed to be independent of 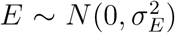 (Paré et al., 2010). In practice, the model often includes other covariates, which are omitted here from notation for simplicity but without loss of generality (Soave and Sun, 2017).

In many GWAS, the interacting *E* may not be measured. Consequently, the working model will be

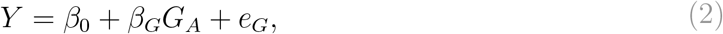

where both *E* and its interaction *G*_*A*_*E* in [1] are latent.

This misspecified working model leads to heteroskedasticity which can be leveraged to indirectly test for the latent interaction. More specifically, the *variance* of the new random error *e*_*G*_ has the following form,

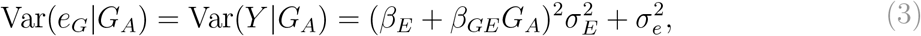

which depends on *G*_*A*_ if *β*_*GE*_*≠* 0. For this reason, various vQTL methods based on Levene-type tests (Paré et al., 2010; Soave and Sun, 2017) or quantile regression method (Miao et al., 2022) have been proposed to identify latent interactions and prioritize genetic variants for follow-up analyses.

### vQTL Approach Does Not Work for Binary Traits

To indirectly test if *β*_*GE*_ = 0 for binary traits, it might seem intuitive to extend the vQTL approach used for quantitative traits. However, the vQTL framework is not applicable to binary traits due to a fundamental difference: unlike a quantitative trait, the variance of a binary trait is inherently determined by its mean,

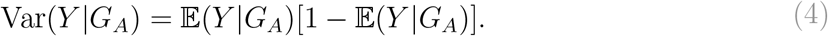

Thus, Var(*Y* |*G*_*A*_) does not yield additional information pertinent to the latent interaction effect. The over-dispersion approach, unfortunately, is not applicable either, as the overdispersion parameter cannot be identified when *Y* is binary (Hinde and Demétrio, 1998).

### Regression Models for Binary Traits

When the trait of interest *Y* is binary, the standard linear association model [1] is replaced by the following generalized linear model (GLM) (Nelder and Wedderburn, 1972),

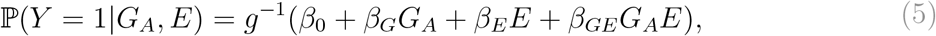

where *G*^*−*1^ refers to the inverse of the GLM link function. Depending on whether a logistic or probit regression is used, *G*^*−*1^ corresponds to the CDF of the standard logistic or normal distribution, respectively.

An equivalent parametrization of model [5] above is through the latent regression formulation (Cramer, 2003),

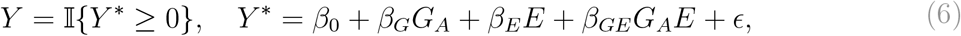

where *Y*\* is latent; the importance of this latent model formulation will be apparent later. Similar to model [1], *G a*nd *E a*re assumed to be independent of each other, and *ϵ i*s assumed to be independent of *G a*nd *E*. But, the error term *ϵ h*ere has a known distribution that is symmetric around zero with CDF *F*_*ϵ*_, whereas the error term *e i*n model [1] typically assumes a normal distribution with an unknown variance 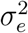. For example, *ϵ c*an follow either the standard logistic distribution or the standard normal distribution, corresponding to a logistic or probit regression model through the GLM formulation in [5].

Given the observed values of *G*_*A*_ and *E*, the conditional probability of being a case (i.e. *Y =* 1) is,

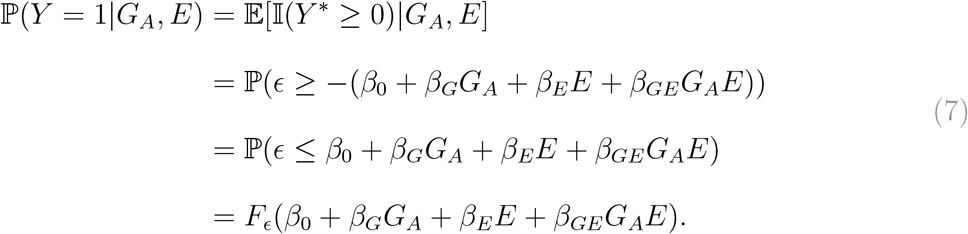

The consequence of missing *E a*nd its interaction in this model will be examined in greater detail in the next section, followed by the proposed method to detect the latent interaction.

## 3. Methods

### Latent Interaction Test Based on Non-Additive Effect for Binary Traits

Assume now the environmental variable *E*, thus also *G*_*A*_*E*, in model [7] is latent, the probability of being a case is now

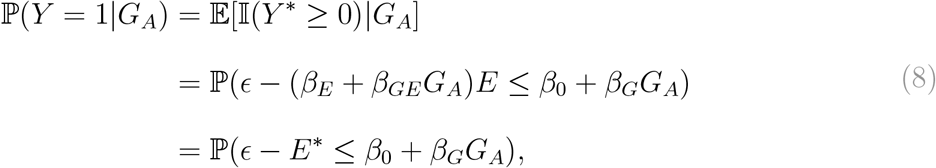

where *E*\* = (*β*_*E*_ + *β*_*GE*_*G*_*A*_)*E*. It is then obvious that *G*_*A*_ and *ϵ − E*\* are independent of each other if and only if *β*_*GE*_ = 0.

To simplify the presentation, we make Assumption 1 without the loss of generality.

#### Assumption 1

*The conditional distribution of ϵ−E*\* *given G*_*A*_ *is in a certain location-scale family. So ϵ*\* = (*ϵ − E*\*)/*SD*(ϵ *− E*\*|*G*_*A*_) *has a completely specified CDF F*_*ϵ*_*∗ that does not depend on G*_*A*_.

#### Remark 1

*When* 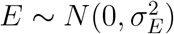, *Assumption 1 will often not hold unless F*_*ϵ*_ *is the standard normal CDF. However, for commonly used models such as logistic regression, Assumption 1 holds approximately due to the close relationship between the standard normal and logistic distributions (Chambers and Cox, 1967). Assumption 1 is only used to simplify the presentation in the rest of this paper; the proposed method remains valid without this assumption*.

If *β*_*GE*_ = 0 and we define *c* = SD(*ϵ − E*\*), Assumption 1 implies that *ϵ*\* = (*ϵ − E*\*)/*c* has a completely specified distribution 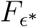, and hence Equation [8] becomes:

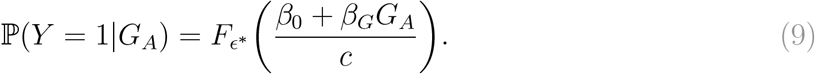

In other words, fitting a binary model with link function 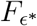 can correctly recover all regression coefficients up to a positive scaling, hence the testing of *β*_*G*_ = 0 is not affected.

When *β*_*GE*_*≠* 0, the variable *ϵ − E*\* will depend on *G*_*A*_ through its standard deviation (SD):

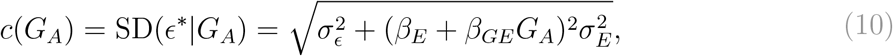

which implies:

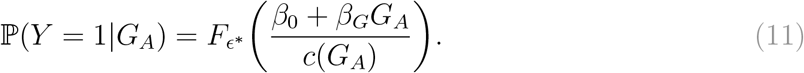

This model is no longer linear on *G*_*A*_, but since *G*_*A*_ only has values of 0, 1 or 2, model [11] is saturated. This implies model [11] can always be fully parameterized with three parameters without the problem of model-misspecification. In particular, model [11] can be rewritten as:

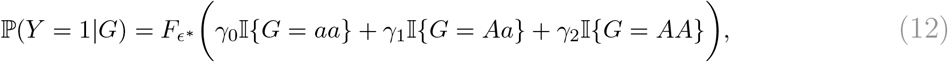

with the parameters *γ*_0_, *γ*_1_ and *γ*_2_ defined as

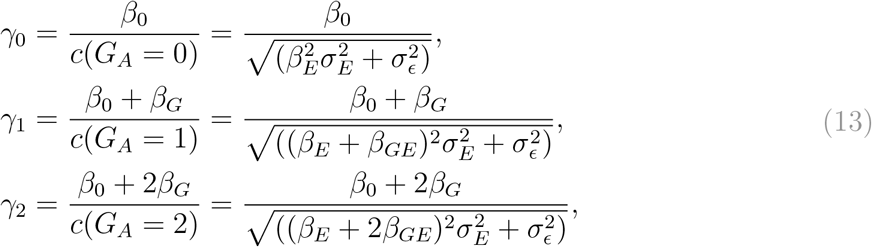

where *σ*_*ϵ*_ is 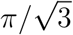 for logistic regression and 1 for probit regression.

#### Remark 2

*In practice, since it is difficult to explicitly know the distribution of ϵ*\* *and hence to fit the corresponding binary model [12], it is easier to fit the binary model with the original link function F*_*ϵ*_. *Since Equation [12] is a saturated model, using a different link function will not introduce any problem of model-inadequacy*.

Define the non-additive effect as:

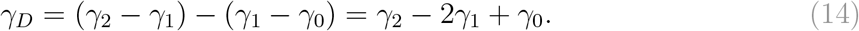

It is clear that if *β*_*GE*_ = 0, the working model [12] is additive with *γ*_*D*_ = 0. If there is latent *G*xE in the original model (i.e. *β*_*GE*_*≠* 0), then an extra non-additive effect *γ*_*D*_ will generally be created by the latent interaction, even when the original model [7] has only an additive effect (*β*_*G*_, *G*_*A*_). Therefore, analogous to using vQTLs to detect *GxE* effect in the analysis of quantitative traits, the *GxE* effect in the analysis of binary traits can be indirectly detected from testing the non-additive effect *γ*_*D*_, which is elsewhere termed the dominance effect.

### Equivalent Parameterization of Genotypic Models

From Equations [11] and [12], it can be noticed that the working model can always be written as a binary regression model with the genotypic encoding in Equation [12]. Since the model is saturated, there are many equivalent re-parametrizations of this genotypic model with three regression parameters. To assess how much non-additive genetic variation is created by the latent *GxE* compared to the additive genetic variation, we consider the Fisher orthogonal re-parametrization of this model under the Hardy-Weinberg equilibrium (HWE) assumption of the SNP:

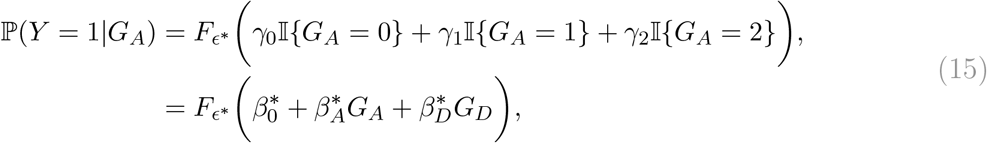

where *G*_*A*_ = (0, 1, 2) and *G*_*D*_ = (*−p*/*q*, 1, *−q*/*p*) for the genotypes (aa, Aa, AA). Given a vector of three genotypic effects ***γ*** = (*γ*_0_, *γ*_1_, *γ*_2_)^*T*^ for the genotypes (*aa, Aa, AA*), the parameters 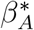 and 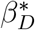 in Equation [15] can be computed as:

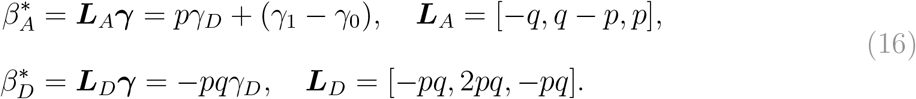

This Fisher orthogonal encoding ensures that the two variables *G*_*A*_ and *G*_*D*_ are uncorrelated (Palmer et al., 2023), and therefore the proportion of genetic effect explained by the non-additive component (on the latent *Y*\*) is

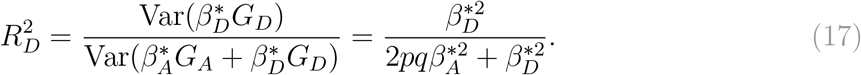

To illustrate the non-additive effect *γ*_*D*_ introduced by the latent *GxE*, we show the contours of *γ*_*D*_ under different parameter settings in Equation [5], for a probit regression model, in Figure [1](A-B). Although the original model [5] only contains the additive component *G*_*A*_, if there exists a latent interaction (i.e. *β*_*GE*_*≠* 0), a non-negligible *γ*_*D*_ is induced for most of the parameter settings. In Figure [1](C-D), we compute the corresponding 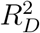 as defined in Equation [17] for the same sets of parameters. In most settings, the induced non-additive component comprises a moderate proportion of the genetic variation; however, when the minor allele of the SNP has a protective effect (*β*_*G*_ *<* 0) for the binary trait, the non-additive proportion is particularly large.

**Figure 1:**
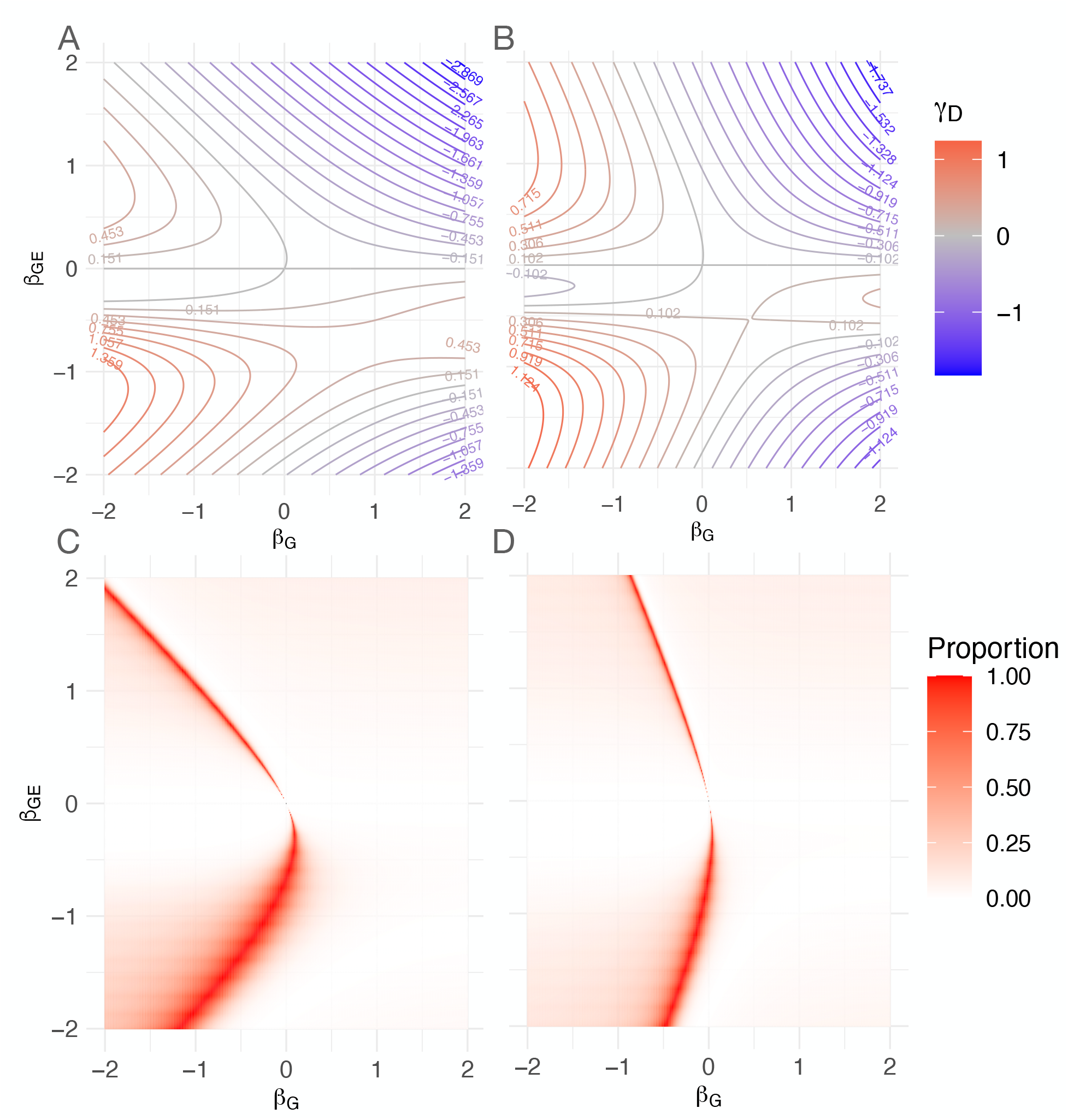
(A-B) show contours of γ_D_ and (C-D) show heat-maps of the non-additive proportion of genetic variation 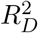, at different β_G_ and β_GE_. The parameter β_E_ = 0.5 and E ∼ N (0, 1). The MAF is set to p = 0.3 in (C-D). The underlying model is assumed to be probit. The prevalence of the binary trait Y is 0.1 on the left column and 0.3 on the right column.

### Wald Test

Although the choice of test does not affect the validity of the proposed method, we adopt the Wald test to be consistent with the current practice of GWAS. Let 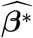 denotes the maximum likelihood estimate (MLE) of the vector of regression parameter 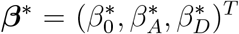 in model [15], *L ∈* R*d×*3 denotes the constraint matrix with *d* linear independent rows for the null hypothesis *H*_0_ : *L****β***\* = **0**. The Wald test uses the test statistics:

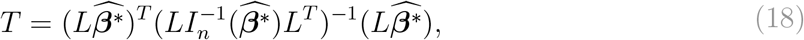

where 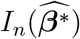 denotes the Fisher information matrix evaluated at the MLE. Under the null hypothesis, the test statistics *T* asymptotically follows a chi-square distribution with *d* degrees of freedom as the sample size *n* grows. To indirectly detect the latent interaction effect *β*_*GE*_, we test the non-additive effect 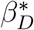 of the SNP, which corresponds to *L* = [0, 0, 1] *∈* R1*×*3. Since 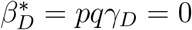 whenever *β*_*GE*_ = 0, the proposed non-additive test will have the correct test size.

In traditional GWAS, the testing of SNPs is typically based on their additive main effects, while omitting possible non-additive effects (Palmer et al., 2023). This additive-only approach corresponds to *L* = [0, 1, 0] *∈* R1*×*3 in the Wald test. Since the latent *GxE* for binary trait (*β*_*GE*_) induces a non-additive genetic effect *γ*_*D*_, or equivalently, 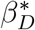 in the working model, we propose a joint test of the hypothesis 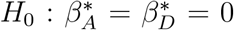 in model [15], in order to detect the latent interaction *β*_*GE*_ together with the main effect *β*_*G*_ in Equation [5]. This uses a constraint matrix *L ∈* R2*×*3 that specifies the null hypothesis 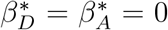, and the Wald test has two degrees of freedom, in contrast to the one degree of freedom test that only considers the additive effect 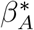. We emphasize that the proposed joint test is not restricted to the Fisher orthogonal encoding in Equation [15]. In fact, the two degrees of freedom joint test can be equivalently performed using any saturated model that encodes the genotypic effects with three regression parameters. The encoding in Equation [15] is used to simplify the partition of additive and non-additive effects.

## 4. Result

### Type I Error Evaluation

In this section, we will assess the type I error rate of the proposed non-additive test and the joint test, respectively for testing *β*_*GE*_ = 0 and *β*_*G*_ = *β*_*GE*_ = 0.

To assess the type I error rate of the proposed non-additive test that detects the latent *GxE* effect *β*_*GE*_ based on the non-additive effect 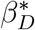, we simulate *n* = 100, 000 independent individuals under the logistic model in Equation [6] under the null hypothesis that *β*_*GE*_ = 0.

We fix *β*_0_ = *−*1 and *β*_*G*_ = 0.5, and independently simulate *E ∼ N*(0, 1) and *G* under the HWE. We consider the MAF of *G* being *{*0.1, 0.3, 0.5*}* and *β*_*E*_ being *{*0, 1*}*. For each of the six settings, the p-values of the proposed test are computed from *B* = 100, 000 independent replications. To also assess the type I error rate of the proposed joint test that accounts for the latent *GxE β*_*GE*_ together with the main effect *β*_*G*_, we further fix the *β*_*G*_ = 0 and obtain the p-values of the proposed joint test from the same settings above. As shown in Table [1], the proposed non-additive and joint tests both have well-controlled type I error rates across different parameter settings. The histograms of the p-values of the non-additive test and the joint test are provided in the supplement (Figures S1-S2), where the distributions of p-values are shown to be close to Unif[0, 1] in all settings.

**Table 1:** Empirical type I error rates of the proposed non-additive test (above) and the joint test (below) for each choice of βE, MAF and significance level α. The rates are computed using B = 100, 000 independent replications, each with n = 100, 000 simulated individuals.

| $\beta_E$ | $\alpha=0.05$ | | | $\alpha=0.005$ | | | $\alpha=0.0005$ | | |
| --- | --- | --- | --- | --- | --- | --- | --- | --- | --- |
|  | MAF=0.1 | 0.3 | 0.5 | 0.1 | 0.3 | 0.5 | 0.1 | 0.3 | 0.5 |
| 0 | 0.0502 | 0.0509 | 0.0517 | 0.00485 | 0.00487 | 0.00512 | 0.00057 | 0.00050 | 0.00056 |
| 1 | 0.0503 | 0.0523 | 0.0520 | 0.00528 | 0.00568 | 0.00575 | 0.00048 | 0.00053 | 0.00050 |
|  | MAF=0.1 | 0.3 | 0.5 | 0.1 | 0.3 | 0.5 | 0.1 | 0.3 | 0.5 |
| 0 | 0.0506 | 0.0498 | 0.0509 | 0.00462 | 0.00541 | 0.00530 | 0.00040 | 0.00053 | 0.00060 |
| 1 | 0.0506 | 0.0504 | 0.0502 | 0.00520 | 0.00485 | 0.00488 | 0.00046 | 0.00052 | 0.00054 |

The p-values and the empirical type I error rates of the proposed non-additive and joint tests above are obtained under a *theoretical* null hypothesis, in which the traits were directly generated from a null model in which the hypothesis *β*_*G*_ = *β*_*GE*_ = 0 is true. As discussed in Zhang and Sun (2019), another way to assess the test size of a method is through the *empirical* null hypothesis, in which the traits are generated from an alternative model, but are then randomly permuted before being tested. To further assess the type I error rate of the proposed tests under the empirical null hypothesis, we use data from the UK Biobank (UKB) (Bycroft et al., 2018) to implement a GWAS in a randomly permuted binary trait (self-reported) high cholesterol (Data-Field 20002; Coding 1473), collected at the baseline. The details of the GWAS procedures are the same as those described later in the next section. The genomic-control (GC) *λ* of the p-values of this permuted GWAS is computed to be 1.004 for the non-additive and 0.996 for the joint test (Devlin et al., 2001). The histogram and the QQ plot of these p-values are displayed in the supplementary material (Figure S3).

### Power Comparison

In this section, we provide a detailed assessment of the powers of the proposed indirect test of the latent interaction *β*_*GE*_, and the powers of the proposed joint test that simultaneously detects the main effect *β*_*G*_ and the latent interaction effect *β*_*GE*_. To simplify the power computation, we assume a probit model in Equation [5] as the true model, which satisfies Assumption 1. The SNP *G* is generated with MAF = 0.3 under the assumption of HWE. The latent environmental variable *E* follows *N*(0, 1) with an effect *β*_*E*_ = *−*0.5, 0 and 0.5. The genetic effect *β*_*G*_ and interaction effect *β*_*GE*_ range from -1 to 1, and the sample size is set to *n* = 30,000, 300,000 and 800,000. The intercept *β*_0_ is set for a prevalence rate of 10 percent.

Since the probit regression model is assumed, the asymptotic power of the Wald test can be computed analytically for each case. First, we compute the corresponding values of 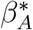 and 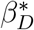 based on the values of *β*_*G*_ and *β*_*GE*_, using Equations [13] and [16]. Second, we compute the non-centrality parameter of the Wald test statistic *T* as

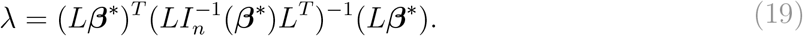

Finally, the asymptotic power of the Wald test is computed using the non-centrality parameter λ as:

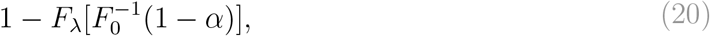

where α is set to the genome-wide significance level 5 *×* 10^*−*8^ (Dudbridge and Gusnanto, 2008); *F*_*λ*_ denotes the CDF of the non-central Chi-square distribution with *d* degrees of freedom and non-centrality parameter *λ*, and 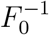 denotes the inverse CDF of the central Chi-square distribution with *d* degrees of freedom.

Figure [2] show the power of the proposed non-additive test of *β*_*GE*_ and the proposed joint test of *β*_*G*_ and *β*_*GE*_, when the sample size *n* = 300,000 or *n* = 30,000. The conclusions from the other settings of sample size are similar and included in the supplementary material. As shown in the figure, both the proposed non-additive and joint tests tend to have higher power when *β*_*GE*_ and *β*_*E*_ have opposite signs, which happens when the environmental variable has opposite effects dependent on the dosage of the minor allele of the SNP. In most cases, when either *β*_*G*_ or *β*_*GE*_ is away from 0, the proposed joint test has power close to 1 to detect the genetic signal. Yet, for certain values of *β*_*G*_ and *β*_*GE*_ that deviate significantly from 0, the joint test exhibits limited power to detect them. This occurs when the values of *β*_*G*_ and *β*_*GE*_ lead to both *β*_*D*_ and *β*_*A*_ in Equation [15] being near zero. As illustrated in the supplement (Figures S5-S7), these instances become less frequent as the sample size *n* increases.

**Figure 2:**
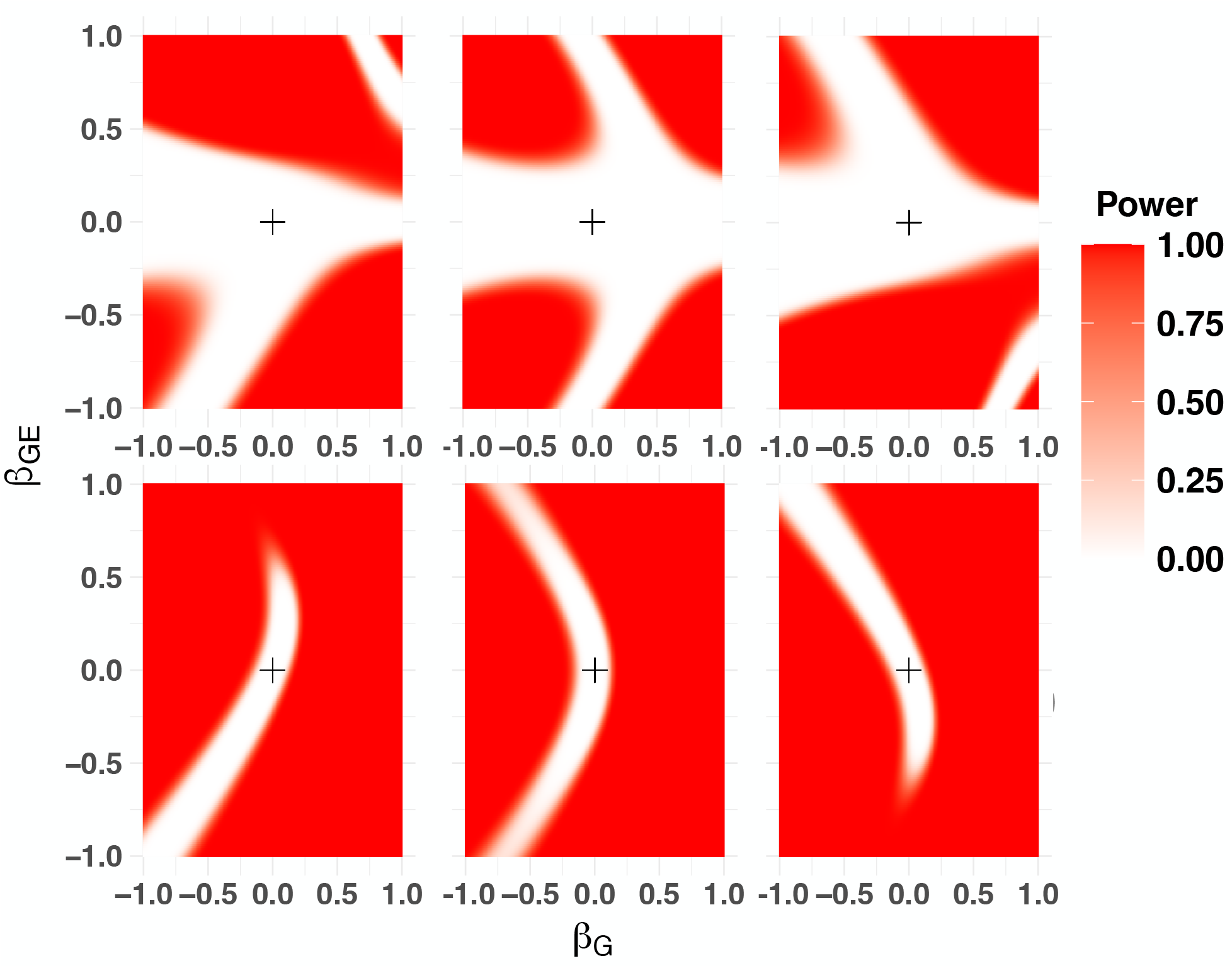
Power of the proposed tests: The power for the proposed non-additive test based on βD is shown in the first row, and the power for the proposed joint test of βGE and βG is shown in the second row. The size of βE is respectively set to ™0.5 (left), 0 (center) and 0.5 (right). The sample size is n = 300,000 on the first row and 30,000 on the second row. The origin in each figure is shown with the cross.

### UKB Application

We illustrated the usage of the proposed non-additive (indirect) test and its subsequent (2-df) joint test. We achieved this by a GWAS on UKB of the binary trait (self-reported) high cholesterol (Data-Field 20002; Coding 1473) (Sudlow et al., 2015; Bycroft et al., 2018).

We selected genotyped SNPs with MAF greater than 0.01, HWE p-values greater than 1e-50 and SNP call rates greater than 0.8. This resulted in 626,164 autosomal SNPs being analyzed. To avoid the potential bias from ancestry, we restricted our analysis to unrelated self-reported British participants with ancestries further confirmed by the PC constructed from genetic data (Data-Field 22006). The related individuals were filtered out based on the kinship coefficients (Data-Field 22021), and we further filtered out individuals with genotype missing rates higher than 0.2. The final sample consists of 276,658 approximately unrelated individuals. The prevalence rate of the trait in the final sample is 0.121 (0.151 in males, 0.095 in females).

We then used logistic regression to analyze the genetic association between each SNP and the binary trait (high cholesterol), accounting for covariate effects of age (Data-Field 21022), sex (Data-Field 31) and first four principal components (PC) constructed from genetic data. We carried out the GWAS using both the (2-df) joint test and the (1-df) non-additive test. The two GWAS results are displayed in Figure [3]. As reflected in the Miami plot, we identified several SNPs with genome-wide significant association with high cholesterol using the proposed joint test (GC *λ* = 1.074). Among these SNPs, the non-additive test (GC *λ* = 1.007) flagged 4 SNPs with genome-wide significant non-additive effects for follow-up studies of latent *GxE* effects, with the top rs7412 (p-value = 1.640e-19) in *APOE*. The QQ plots and histograms of the two GWAS can be found in the supplementary material (Figure S4).

**Figure 3:**
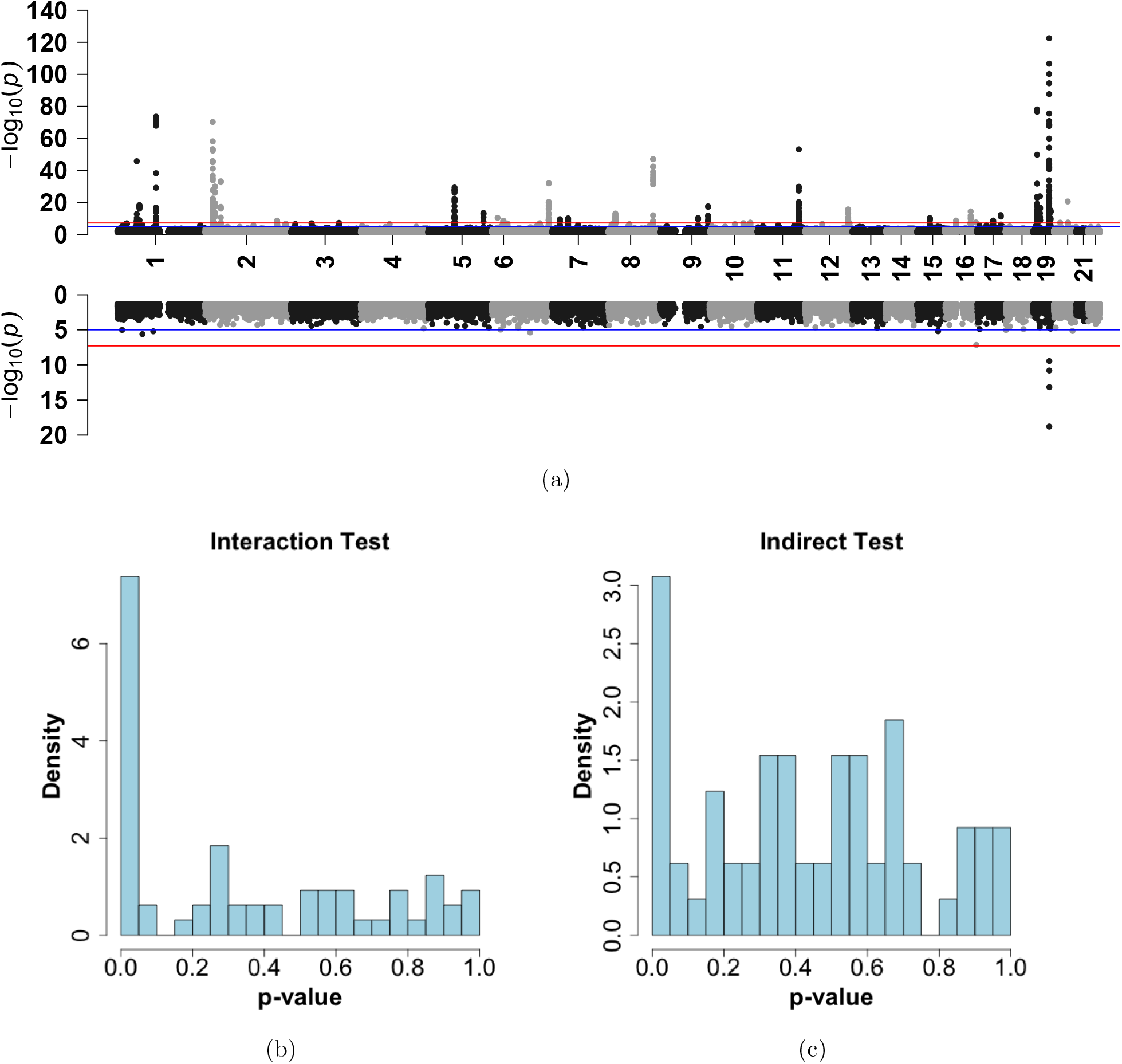
(a): GWAS result using the proposed 2-df joint test (upper) and the indirect non-additive test (bottom). The red line denotes the genome-wide significance level of 5e-8. (b-c): The histogram of p-values for the 65 selected SNPs to test either the interaction effect with rs7412 (b) or the non-additive effect (c).

The Manhattan plot for the traditional GWAS based on the additive test is also included in the supplementary material (Figure S8).

The latent interactions suggested by the non-additive test are not unexpected, given the well-established literature on the haplotype effects of *APOE* on cholesterol levels (Murdoch et al., 2007), which can be viewed as an interaction between nearby SNPs (Ken-Dror et al., 2013). To further investigate the latent interactions with nearby SNPs, we selected 65 SNPs with *D*^*′*^ larger than 0.2 within 10,000 kb of rs7412, and then performed the GxG interaction analysis between those SNPs and rs7412. The LD information including *D*^*′*^, *r*^2^ as well as the physical position of these SNPs were obtained using the tool LDlink (Machiela and Chanock, 2015), with the genome build GRCh37 and super-population of all the European groups (CEU, TSI, FIN, GBR and IBS). The histograms of p-values for the interaction tests and for the proposed indirect tests of the selected 65 SNPs are provided in Figure [3]. Indeed, we found 11 SNPs having interaction with rs7412 at the significance level of 0.05 after Bonferroni correction; 5 of the 11 SNPs also have p-values less than 0.05 using the proposed non-additive test based on the non-additive effect. The SNP with the smallest p-value of the interaction test (1.713e-07) is rs7254892, which is mapped to *NECTIN2*. This SNP has *D*^*′*^ = 1 with rs429358, which in combination with rs7412 defines the classic *APOE* haplotypes (*ϵ*_2_, *ϵ*_3_, *ϵ*_4_) (Murdoch et al., 2007). The detailed results for the 11 SNPs are provided in Table [2].

**Table 2:** Summary of characteristics of the 11 SNPs identified through the interaction analysis; including minor allele frequencies, linkage disequilibrium measures (D’ and R^2^), distance (in BP) to rs7412, and association test p-values (Indirect and Interaction) along with estimated regression coefficients 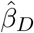and 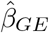. As a comparison, the D’ to rs429358 is also shown in the parenthesis.

| Rank | SNP | MAF | $D'$ | $R^2$ | Distance | P:Indirect | P:Interaction | $\hat{\beta}_D$ | $\hat{\beta}_{GE}$ |
| --- | --- | --- | --- | --- | --- | --- | --- | --- | --- |
| 1 | rs7254892 | 0.031 | 0.931 (1.000) | 0.413 | -22483 | 3.723e-03 | 1.713e-07 | -0.0184 | 0.04826 |
| 2 | rs141622900 | 0.045 | 0.953 (1.000) | 0.636 | 14713 | 6.685e-14 | 1.960e-07 | -0.0474 | 0.06425 |
| 3 | rs34954997 | 0.219 | 1.000 (0.984) | 0.239 | 5559 | 3.402e-01 | 2.733e-07 | -0.0060 | 0.05764 |
| 4 | rs483082 | 0.219 | 1.000 (0.984) | 0.239 | 4099 | 3.166e-01 | 4.940e-07 | -0.0063 | 0.05684 |
| 5 | rs405509 | 0.484 | 1.000 (0.602) | 0.063 | -3243 | 3.849e-01 | 3.467e-06 | -0.0052 | -0.04932 |
| 6 | rs440446 | 0.363 | 1.000 (0.982) | 0.038 | -2912 | 9.960e-01 | 4.849e-06 | 0.0000 | -0.04909 |
| 7 | rs75627662 | 0.186 | 1.000 (0.748) | 0.293 | 1497 | 3.514e-02 | 6.094e-06 | -0.0134 | 0.05253 |
| 8 | rs72654473 | 0.093 | 1.000 (0.182) | 0.648 | 2320 | 3.613e-10 | 1.337e-05 | -0.0425 | 0.06595 |
| 9 | rs439401 | 0.381 | 1.000 (0.983) | 0.041 | 2372 | 6.640e-01 | 2.835e-05 | 0.0026 | -0.04502 |
| 10 | rs584007 | 0.378 | 1.000 (0.983) | 0.041 | 4399 | 8.491e-01 | 8.011e-05 | -0.0012 | -0.04235 |
| 11 | rs445925 | 0.094 | 1.000 (0.190) | 0.641 | 3561 | 1.593e-11 | 1.825e-04 | -0.0419 | 0.06027 |

To confirm that accounting for the interaction with rs7412 can explain the non-additive effects flagged earlier by the proposed non-additive test, we performed the non-additive test on the 65 selected SNPs both with or without considering the interaction with rs7412. When the interaction is not considered, we found 10 SNPs with p-values from the non-additive test less than 0.05, and 3 SNPs with p-values less than 5e-8. After accounting for the interaction effect, the (-log10) p-values, and the magnitudes of the estimated non-additive effects, are shrunk for 8 of the 10 SNPs, as summarized in Figure [4]. In particular, none of the 10 SNP has a genome-wide significant p-value of the non-additive test after their interactions with rs7412 are accounted for.

**Figure 4:**
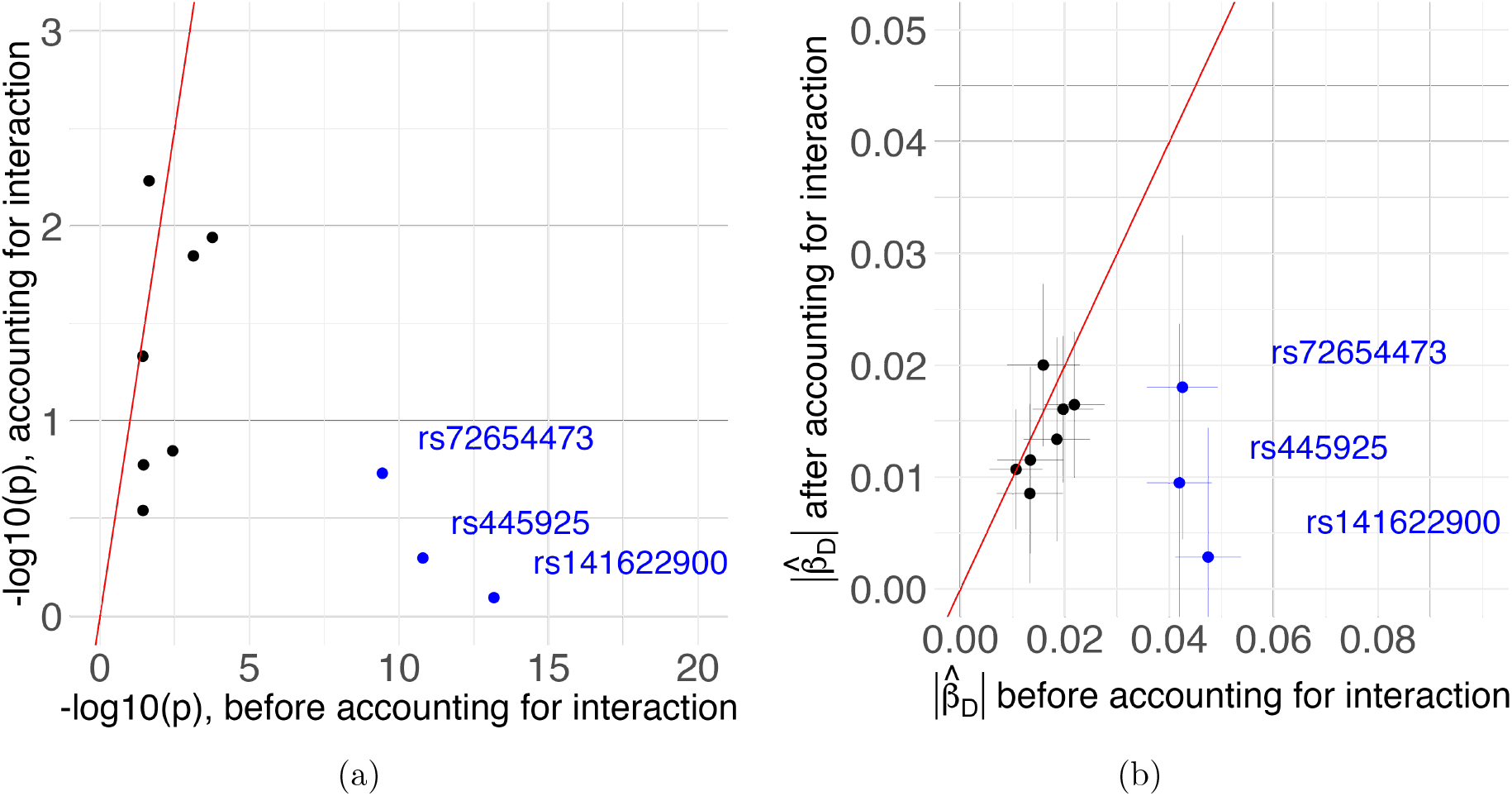
The (-log10) p-values of the non-additive test (a) and the absolute values of estimated non-additive effects (b) before (x-axis) and after (y-axis) accounting for the interaction with rs7412, for the ten SNPs with p-values of non-additive test less than 0.05. The three SNPs with genomewide significant p-values of the non-additive test before accounting the interaction are highlighted in blue. The red line is the line of y = x. The radius of each cross in (b) denotes the standard error of the non-additive effect estimate.

## 5. Discussion

Using heteroskedasticity to indirectly test for a *GxE* is well-established in the analysis of quantitative traits, and has led to many scientific insights over the human genome. However, none of the existing approaches of indirect testing could be applied when the trait of interest is binary. In this paper, we propose the first approach to indirectly test for a *GxE* in the analysis of binary traits, based on the non-additive effect of the genetic variant, and subsequently propose a joint test that could account for the latent *GxE* in the binary trait GWAS. We have applied this method both in the simulation studies and the analysis of the binary trait hypertension in the UKB data, and found promising SNPs with supporting evidence from the existing literature.

It has been suggested in the literature that non-additive genetic effects do not explain as much variability as the additive effects in most human traits (Palmer et al., 2023), supported by the weak dominance signals identified from the dominance GWAS scan. Furthermore, Iles (2010) has shown for binary traits that the non-additive signals are breaking down more rapidly as the linkage disequilibrium breaks down, which has been viewed as another reason to prefer the use of the additive-only model and to ignore the non-additive component in the analysis. Although non-additive signals may not be as prevalent as the additive signals across the genome in a marginal scan, our works have suggested that joint testing of the non-additive and the additive effect may uncover many SNPs that could not be identified by the traditional additive test alone. At the same time, our study illustrates that for binary traits, the additive and non-additive effects of SNPs cannot be easily interpreted separately. Therefore, these two effects should be jointly tested and interpreted together as the genetic effect in binary trait GWAS.

Our approach has a similar nature to the approach of Soave et al. (2015) for quantitative traits, where Soave et al. (2015) accounts for latent *GxE* by jointly testing the genetic effects in the location and the scale of the quantitative trait, and our approach accounts for *GxE* by jointly testing the additive and non-additive genetic effects for the binary trait. In Soave et al. (2015), it is emphasized that we cannot conclude whether the heteroskedasticy is caused by the SNP itself, or by a latent *GxE*. Similarly for our approach, a significant non-additive effect could be either due to the biological mechanism of the SNP itself, or its interaction with latent variables. These joint tests provide valuable insights to screen out SNPs for more detailed interaction analysis, but their results should not be over-interpreted.

## Acknowledgements

Ziang Zhang is a trainee of the CANSSI-ONTARIO STAGE (Strategic Training for Advanced Genetic Epidemiology) training program at the University of Toronto.

## Funding

This research was funded by the Natural Sciences and Engineering Research Council of Canada (NSERC, RGPIN-04934), the Center for Addition and Mental Health (CAMH) Discovery Fund Seed Funding, and the University of Toronto Data Sciences Institute (DSI) Catalyst Grant.

## Data Availability Statement

This research has been conducted using the UK Biobank Resource under Application Number 64875. Data are available at www.ukbiobank.ac.uk/ with the permission of UK Biobank.

## Ethics Statement

The ethics approval of UK Biobank has been obtained from the North West Multi-centre Research Ethics Committee (MREC).

## Notes

### Competing Interest Statement

The authors have declared no competing interest.

