## Supplemental Material for "Detecting latent gene-environment interaction when analyzing binary traits"

### Supplementary Material for "Detecting latent gene-environment interaction when analyzing binary traits"

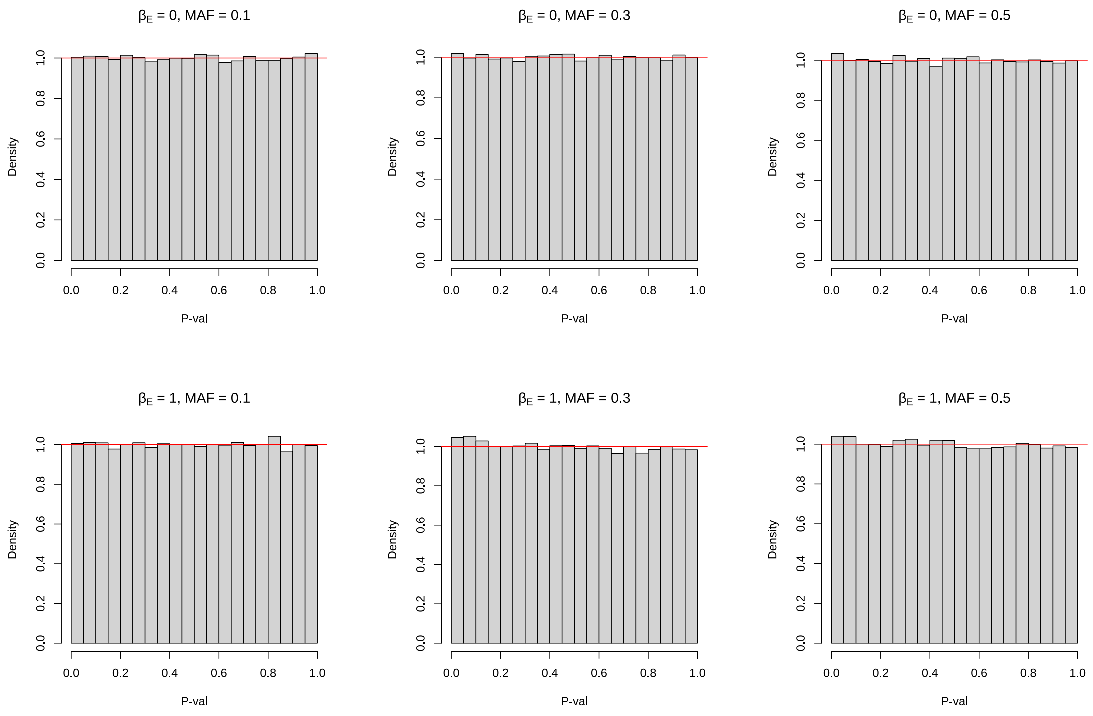


Figure S1: The histograms for the p-values of the proposed indirect test when $\beta_{GE} = 0$, at different settings of the MAF and $\beta_E$. The individuals ($n = 100,000$) are simulated from a logistic regression model with $\beta_G = 0.5$ and $\beta_0 = -1$. The latent variable $E$ follows $N(0, 1)$ independent of the SNP.

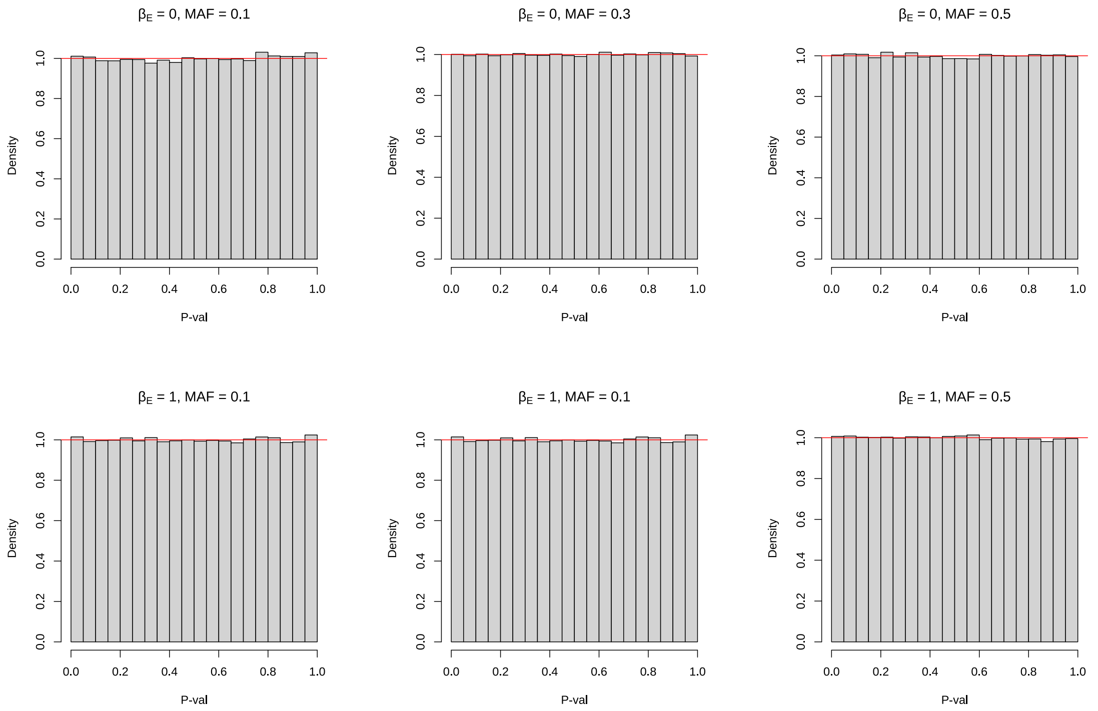


Figure S2: The histograms for the p-values of the proposed joint test when $\beta_{GE} = \beta_G = 0$, at different settings of the MAF and $\beta_E$. The individuals ($n = 100,000$) are simulated from a logistic regression model with $\beta_0 = -1$. The latent variable $E$ follows $N(0,1)$ independent of the SNP.

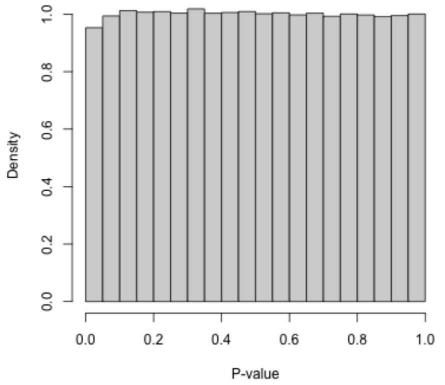


(a) Permuted (indirect) GWAS: Histogram

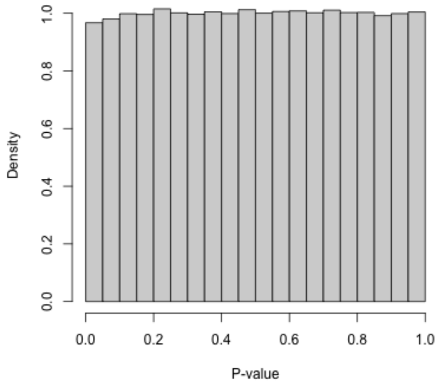


(b) Permuted (joint) GWAS: Histogram

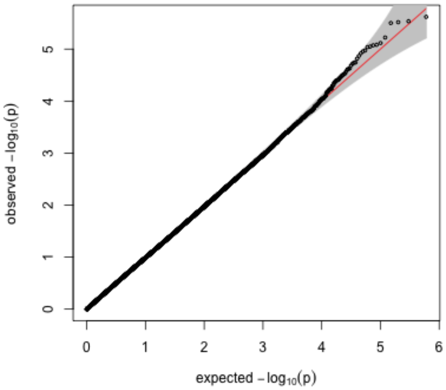


(c) Permuted (indirect) GWAS: QQ-Plot

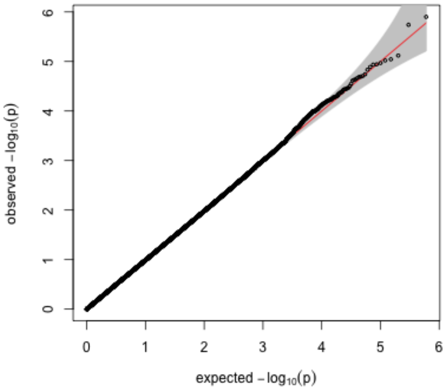


(d) Permuted (joint) GWAS: QQ-Plot

Figure S3: The histograms (a-b) and QQ-plots (c-d) for the GWAS p-values (indirect in the left, joint in the right), for the European population. The binary trait (high cholesterol) has been permuted before the GWAS.

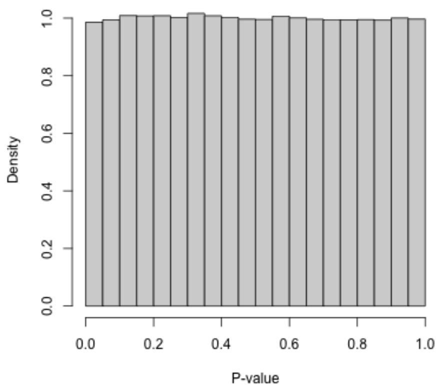


(a) (indirect) GWAS: Histogram

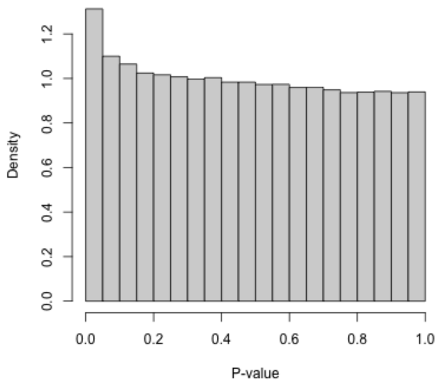


(b) joint) GWAS: Histogram

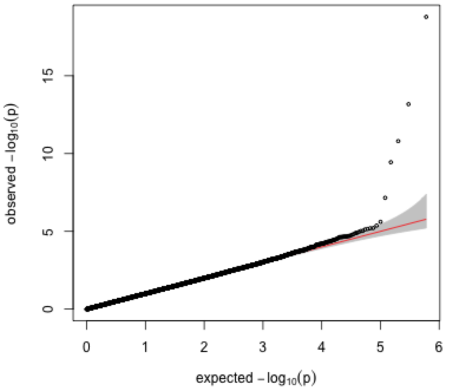


(c) (indirect) GWAS: QQ-Plot

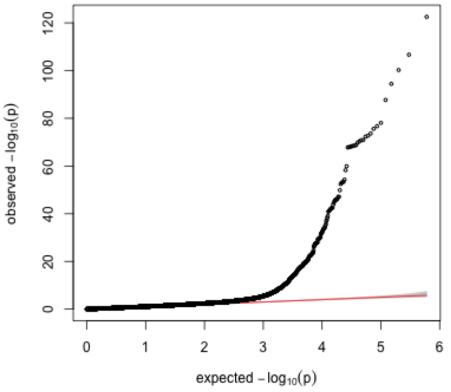


(d) (joint) GWAS: QQ-Plot

Figure S4: The histograms (a-b) and QQ-plots (c-d) for the GWAS p-values (indirect in the left, joint in the right), for the European population.

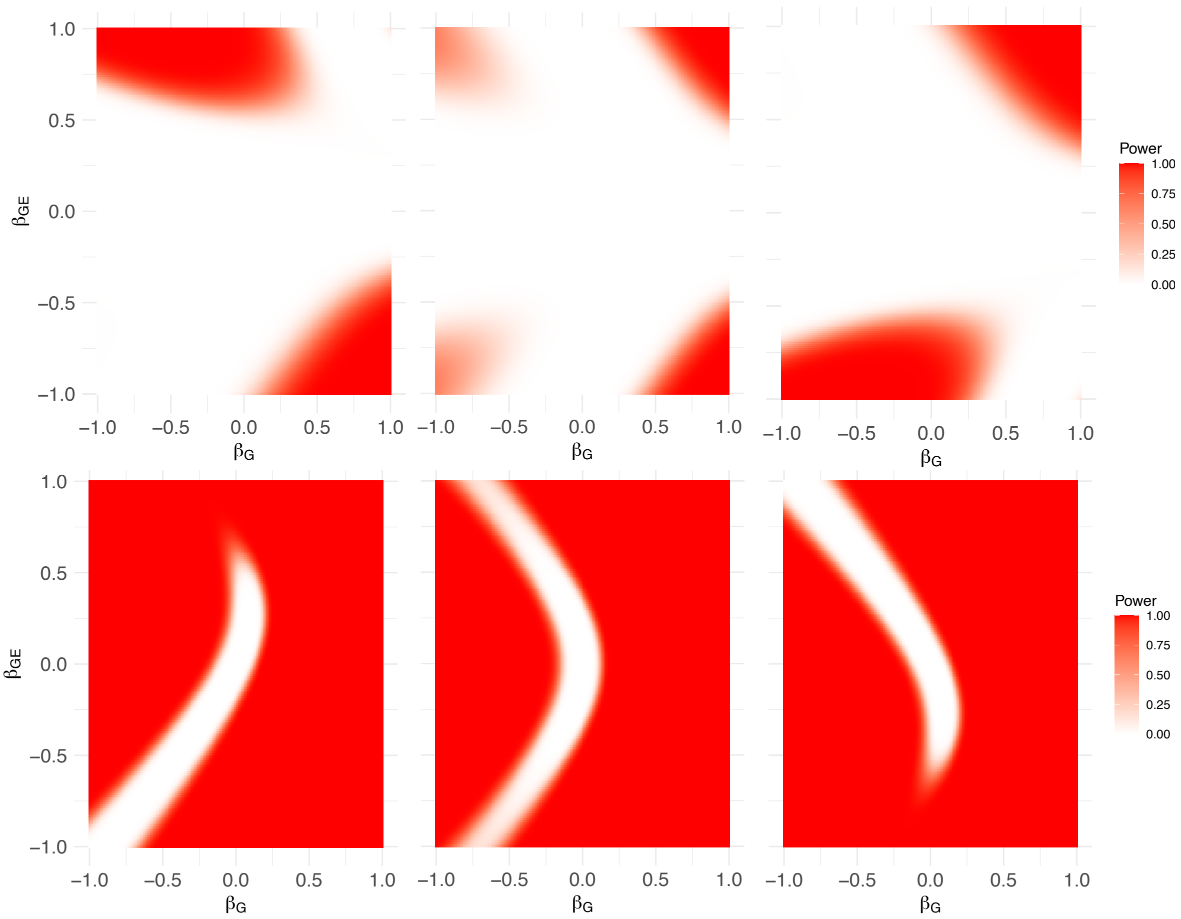


Figure S5: Power of the proposed tests: The power for the proposed indirect test of $\beta_{GE}$ based on the non-additive effect $\beta_D$ is shown in the first row, and the power for the proposed joint test of $\beta_{GE}$ and $\beta_G$ is shown in the second row. The size of $\beta_E$ is respectively set to $-0.5$ (left), $0$ (center) and $0.5$ (right). The sample size is $n = 30,000$.

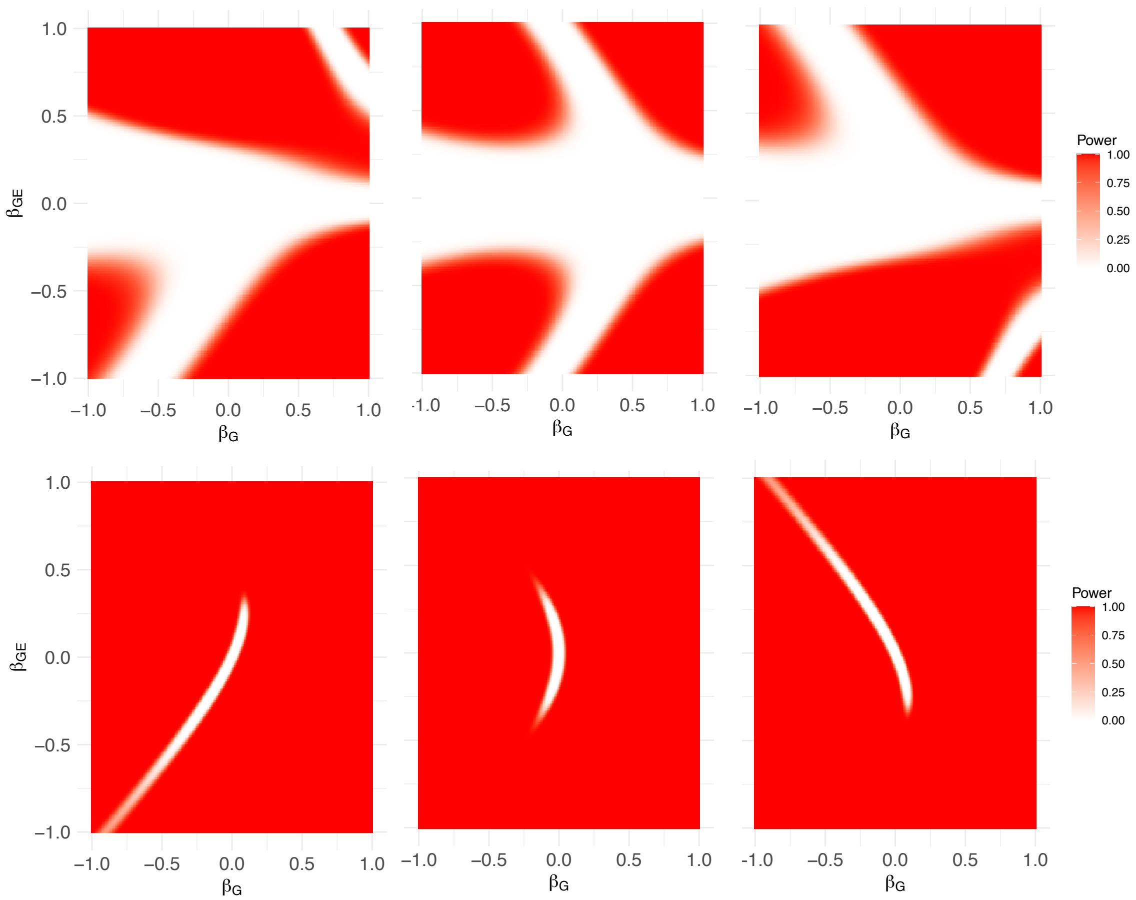


Figure S6: Power of the proposed tests: The power for the proposed non-additive based on $\beta_D$ is shown in the first row, and the power for the proposed joint test of $\beta_{GE}$ and $\beta_G$ is shown in the second row. The size of $\beta_E$ is respectively set to $-0.5$ (left), 0 (center) and 0.5 (right). The sample size is $n$ = 300,000, which is comparable to the size of modern Biobanks.

| Model | Specification | Parameters |
|---|---|---|
| Original model | $\beta_0 + \beta_G G_A + \beta_E E + \beta_{GE} G_A E$<br>$G_A = (0, 1, 2)$ for $(aa, Aa, AA)$ | - |
| Saturated model | $\gamma_0 I(G = aa) + \gamma_1 I(G = Aa) + \gamma_2 I(G = AA)$ | $\gamma_0 = \frac{\beta_0}{\beta_E^2 \sigma_E^2 + 1}$,<br>$\gamma_1 = \frac{\beta_0 + \beta_G}{(\beta_E + \beta_{GE})^2 \sigma_E^2 + 1}$,<br>$\gamma_2 = \frac{\beta_0 + 2\beta_G}{(\beta_E + 2\beta_{GE})^2 \sigma_E^2 + 1}$,<br>$\gamma_D = \gamma_2 - 2\gamma_1 + \gamma_0$<br>(if $\beta_{GE} = 0$ then $\gamma_D = 0$). |
| Fisher orthogonal coding | $\beta_0^* + \beta_A^* G_A + \beta_D^* G_D$<br>$G_A = (0, 1, 2)$ for $(aa, Aa, AA)$<br>$G_D = (-p/q, 1, -q/p)$ for $(aa, Aa, AA)$ | $\beta_0^* = (1 - p^2)\gamma_0 + p^2(2\gamma_1 - \gamma_2)$,<br>$\beta_A^* = -q\gamma_0 - (p - q)\gamma_1 + p\gamma_2$<br>$= p\gamma_D + (\gamma_1 - \gamma_0)$,<br>(if $\beta_{GE} = 0$ then $\beta_A^* = \beta_G$).<br>$\beta_D^* = -pq\gamma_D$ |

Table S1: Different parameterization of the saturated model: The parameters parameterization of $\gamma_0, \gamma_1, \gamma_2$ in terms of $\beta_0, \beta_G, \beta_{GE}$ assumes that the original model is probit. However, as explained in the paper, there will be no model-misspecification issue even when the original model is not probit. The notation $p$ denotes the minor allele frequency and $q = 1 - p$.

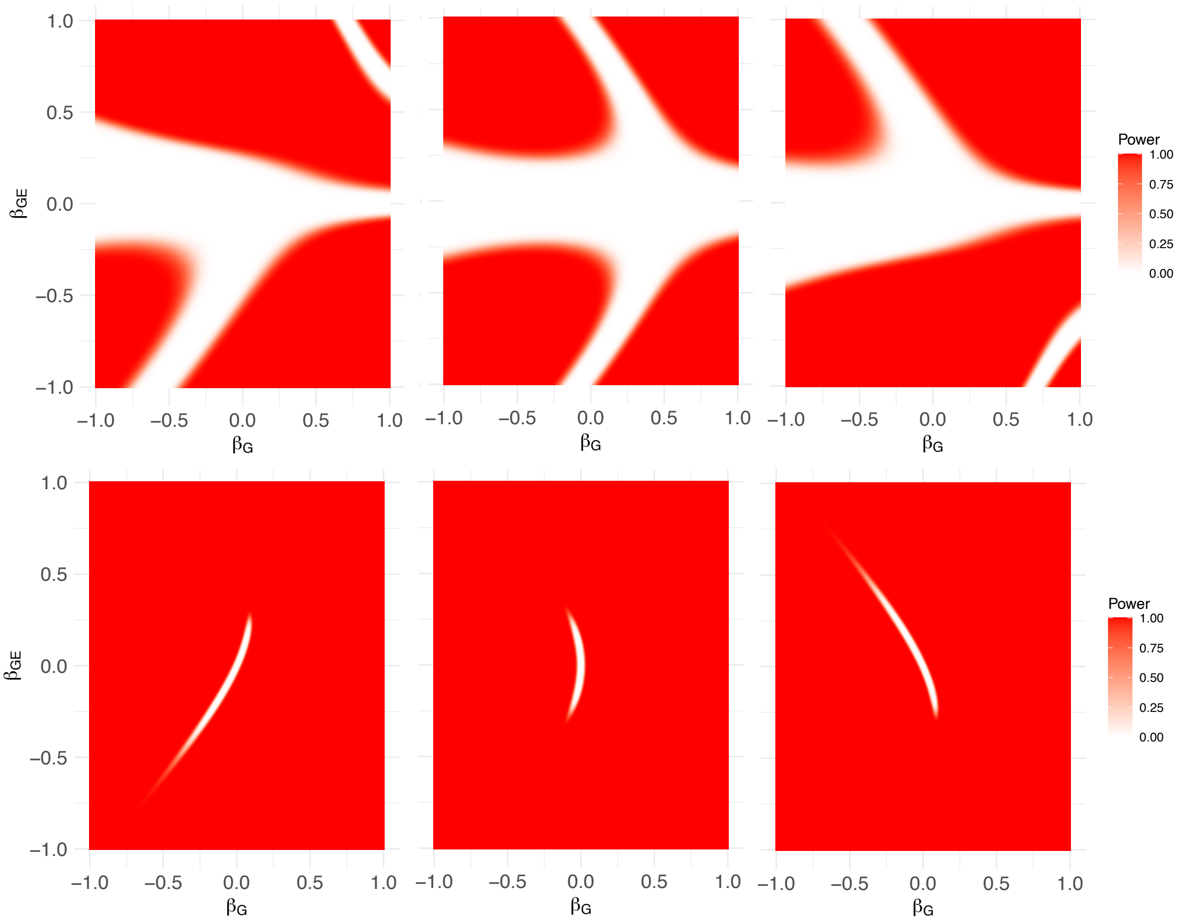


Figure S7: Power of the proposed tests: The power for the proposed indirect test of $\beta_{GE}$ based on the non-additive effect $\beta_D$ is shown in the first row, and the power for the proposed joint test of $\beta_{GE}$ and $\beta_G$ is shown in the second row. The size of $\beta_E$ is respectively set to $-0.5$ (left), 0 (center) and 0.5 (right). The sample size is $n = 800,000$.

Table S2: Summary of characteristics of all the 65 SNPs studied in the interaction analysis; including minor allele frequencies, linkage disequilibrium measures (D' and $R^2$), distance (in BP) to rs7412, and association test p-values (Indirect and Interaction) along with estimated regression coefficients ($\hat{\beta}_D$ and $\hat{\beta}_{GE}$).

| | SNP | MAF | $D'$ | $R^2$ | Distance | p: Indirect | p: Interaction | $\hat{\beta}_D$ | $\hat{\beta}_{GE}$ |
|---|---|---|---|---|---|---|---|---|---|
| 1 | rs7254892 | 0.03 | 0.93 | 0.41 | -22483 | 3.723e-03 | 1.713e-07 | -1.843e-02 | 4.826e-02 |
| 2 | rs141622900 | 0.04 | 0.95 | 0.64 | 14713 | 6.685e-14 | 1.960e-07 | -4.742e-02 | 6.425e-02 |
| 3 | rs34954997 | 0.22 | 1.00 | 0.24 | 5559 | 3.402e-01 | 2.733e-07 | -6.002e-03 | 5.764e-02 |
| 4 | rs483082 | 0.22 | 1.00 | 0.24 | 4099 | 3.166e-01 | 4.940e-07 | -6.291e-03 | 5.684e-02 |
| 5 | rs405509 | 0.48 | 1.00 | 0.06 | -3243 | 3.849e-01 | 3.467e-06 | -5.203e-03 | -4.932e-02 |
| 6 | rs440446 | 0.36 | 1.00 | 0.04 | -2912 | 9.960e-01 | 4.849e-06 | -3.244e-05 | -4.909e-02 |
| 7 | rs75627662 | 0.19 | 1.00 | 0.29 | 1497 | 3.514e-02 | 6.094e-06 | -1.337e-02 | 5.253e-02 |
| 8 | rs72654473 | 0.09 | 1.00 | 0.65 | 2320 | 3.613e-10 | 1.337e-05 | -4.249e-02 | 6.595e-02 |
| 9 | rs439401 | 0.38 | 1.00 | 0.04 | 2372 | 6.640e-01 | 2.835e-05 | 2.638e-03 | -4.502e-02 |
| 10 | rs584007 | 0.38 | 1.00 | 0.04 | 4399 | 8.491e-01 | 8.011e-05 | -1.235e-03 | -4.235e-02 |
| 11 | rs445925 | 0.09 | 1.00 | 0.64 | 3561 | 1.593e-11 | 1.825e-04 | -4.191e-02 | 6.027e-02 |
| 12 | rs62120566 | 0.01 | 0.43 | 0.04 | -214347 | 2.741e-01 | 2.564e-03 | -7.277e-03 | 1.535e-02 |
| 13 | rs7259004 | 0.10 | 0.51 | 0.16 | 20478 | 6.523e-01 | 3.433e-03 | -2.774e-03 | 1.706e-02 |
| 14 | rs405697 | 0.27 | 1.00 | 0.02 | -7388 | 6.944e-01 | 3.688e-03 | 2.566e-03 | -3.079e-02 |
| 15 | rs62119327 | 0.01 | 0.29 | 0.02 | -225206 | 1.582e-01 | 8.136e-03 | -9.523e-03 | 1.322e-02 |
| 16 | rs2965169 | 0.44 | 0.63 | 0.03 | -160923 | 3.917e-01 | 1.229e-02 | -5.213e-03 | 2.014e-02 |
| 17 | rs184017 | 0.21 | 0.36 | 0.03 | -17110 | 3.901e-01 | 1.476e-02 | 5.214e-03 | -1.636e-02 |
| 18 | rs283815 | 0.21 | 0.36 | 0.03 | -21746 | 3.202e-01 | 1.788e-02 | 6.073e-03 | -1.593e-02 |
| 19 | rs59007384 | 0.21 | 0.36 | 0.03 | -15414 | 1.944e-01 | 1.867e-02 | 7.893e-03 | -1.566e-02 |
| 20 | rs157581 | 0.21 | 0.36 | 0.03 | -16365 | 5.273e-01 | 2.259e-02 | 3.600e-03 | -1.525e-02 |
| 21 | rs76560105 | 0.02 | 0.47 | 0.08 | -112880 | 1.745e-04 | 2.285e-02 | -2.179e-02 | 1.148e-02 |
| 22 | rs157582 | 0.21 | 0.36 | 0.03 | -15860 | 3.239e-01 | 2.386e-02 | 6.004e-03 | -1.511e-02 |
| 23 | rs28399637 | 0.32 | 0.66 | 0.01 | -87941 | 5.845e-02 | 2.708e-02 | 1.117e-02 | -2.074e-02 |
| 24 | rs11668861 | 0.46 | 0.79 | 0.03 | -31109 | 4.978e-01 | 2.890e-02 | -4.451e-03 | -2.138e-02 |
| 25 | rs28399654 | 0.03 | 0.45 | 0.09 | -95491 | 7.565e-04 | 7.091e-02 | -1.964e-02 | 9.116e-03 |
| 26 | rs11668327 | 0.16 | 0.42 | 0.06 | -13446 | 2.293e-02 | 7.287e-02 | 1.583e-02 | 1.218e-02 |
| 27 | rs10411821 | 0.33 | 0.29 | 0.01 | 684053 | 6.518e-01 | 1.679e-01 | 2.713e-03 | 1.002e-02 |
| 28 | rs519113 | 0.21 | 0.33 | 0.03 | -35795 | 4.205e-01 | 2.099e-01 | -4.921e-03 | 8.759e-03 |
| 29 | rs2965101 | 0.35 | 0.39 | 0.02 | -174267 | 1.031e-01 | 2.162e-01 | -9.945e-03 | 8.969e-03 |
| 30 | rs8109620 | 0.31 | 0.40 | 0.02 | 250380 | 3.968e-01 | 2.601e-01 | -5.087e-03 | -8.093e-03 |
| 31 | rs10405693 | 0.34 | 0.72 | 0.02 | -85415 | 8.591e-01 | 2.620e-01 | -1.044e-03 | -1.080e-02 |
| 32 | rs6859 | 0.44 | 0.82 | 0.04 | -30045 | 5.918e-01 | 2.631e-01 | -3.184e-03 | -1.053e-02 |

Continued on next page

**Table S2 continued from previous page**

| | SNP | MAF | $D'$ | $R^2$ | Distance | p: Indirect | p: Interaction | $\hat{\beta}_D$ | $\hat{\beta}_{GE}$ |
|---|---|---|---|---|---|---|---|---|---|
| 33 | rs1531517 | 0.05 | 0.36 | 0.11 | -169906 | 8.796e-01 | 2.647e-01 | -1.010e-03 | 6.161e-03 |
| 34 | rs429358 | 0.16 | 1.00 | 0.01 | -138 | 1.988e-01 | 2.842e-01 | 7.619e-03 | -1.079e-02 |
| 35 | rs4420638 | 0.20 | 0.92 | 0.01 | 10867 | 3.210e-01 | 2.927e-01 | 5.483e-03 | -1.025e-02 |
| 36 | rs365653 | 0.09 | 0.37 | 0.09 | -50433 | 5.045e-01 | 3.009e-01 | -4.677e-03 | 6.320e-03 |
| 37 | rs28399653 | 0.03 | 0.45 | 0.09 | -96634 | 5.861e-02 | 3.257e-01 | -1.153e-02 | 5.109e-03 |
| 38 | rs12721109 | 0.02 | 0.76 | 0.16 | 35142 | 3.646e-02 | 3.660e-01 | -1.327e-02 | -5.204e-03 |
| 39 | rs8104483 | 0.29 | 0.73 | 0.01 | -39725 | 5.929e-01 | 3.955e-01 | 3.165e-03 | -7.920e-03 |
| 40 | rs283813 | 0.08 | 0.42 | 0.14 | -22905 | 1.689e-01 | 4.030e-01 | -8.716e-03 | -4.444e-03 |
| 41 | rs11672271 | 0.31 | 0.42 | 0.03 | 207867 | 7.355e-01 | 4.150e-01 | -2.030e-03 | -5.871e-03 |
| 42 | rs16979890 | 0.14 | 0.20 | 0.02 | 654239 | 7.465e-01 | 5.052e-01 | 1.953e-03 | 4.656e-03 |
| 43 | rs4803770 | 0.36 | 1.00 | 0.04 | 15274 | 9.462e-01 | 5.259e-01 | 4.303e-04 | -6.574e-03 |
| 44 | rs8106922 | 0.41 | 1.00 | 0.05 | -10413 | 8.845e-01 | 5.262e-01 | 9.294e-04 | 6.321e-03 |
| 45 | rs71352239 | 0.30 | 0.95 | 0.03 | 17464 | 9.858e-01 | 5.586e-01 | -1.125e-04 | 5.636e-03 |
| 46 | rs12162222 | 0.30 | 0.63 | 0.01 | -63557 | 5.451e-01 | 5.789e-01 | -3.543e-03 | -5.352e-03 |
| 47 | rs62117160 | 0.03 | 0.71 | 0.20 | -179918 | 2.440e-01 | 5.993e-01 | -7.985e-03 | 2.973e-03 |
| 48 | rs3852856 | 0.21 | 0.78 | 0.01 | -50505 | 9.406e-01 | 6.040e-01 | 4.713e-04 | -5.208e-03 |
| 49 | rs16979873 | 0.14 | 0.20 | 0.02 | 622432 | 9.010e-01 | 6.164e-01 | -7.517e-04 | -3.506e-03 |
| 50 | rs35891370 | 0.35 | 0.29 | 0.01 | 107021 | 9.746e-01 | 6.499e-01 | 1.908e-04 | -3.283e-03 |
| 51 | rs59325138 | 0.40 | 1.00 | 0.04 | 4212 | 6.269e-01 | 6.530e-01 | 3.101e-03 | -4.655e-03 |
| 52 | rs769450 | 0.41 | 1.00 | 0.05 | -1635 | 6.064e-01 | 7.116e-01 | 3.079e-03 | -3.888e-03 |
| 53 | rs3745150 | 0.44 | 0.93 | 0.04 | -26320 | 5.988e-01 | 7.711e-01 | -3.358e-03 | -2.973e-03 |
| 54 | rs3178166 | 0.46 | 0.41 | 0.01 | 182091 | 2.892e-01 | 7.807e-01 | 6.357e-03 | -2.088e-03 |
| 55 | rs421812 | 0.29 | 0.78 | 0.02 | -31534 | 6.864e-01 | 7.920e-01 | -2.603e-03 | 2.480e-03 |
| 56 | rs45564734 | 0.02 | 0.26 | 0.02 | -904898 | 3.584e-01 | 8.192e-01 | -4.986e-03 | -1.551e-03 |
| 57 | rs73045960 | 0.02 | 0.37 | 0.04 | 883144 | 3.692e-02 | 8.684e-01 | -1.062e-02 | -8.778e-04 |
| 58 | rs11881756 | 0.10 | 0.22 | 0.03 | -191183 | 4.018e-01 | 8.686e-01 | -5.260e-03 | -9.878e-04 |
| 59 | rs34978331 | 0.23 | 0.24 | 0.01 | -79238 | 2.397e-01 | 8.939e-01 | 7.161e-03 | 9.333e-04 |
| 60 | rs385982 | 0.33 | 0.76 | 0.02 | -32397 | 6.673e-01 | 8.981e-01 | -2.768e-03 | -1.212e-03 |
| 61 | rs369599 | 0.29 | 0.78 | 0.02 | -32743 | 5.606e-01 | 9.186e-01 | -3.742e-03 | -9.678e-04 |
| 62 | rs3837923 | 0.29 | 0.78 | 0.02 | -32513 | 5.151e-01 | 9.483e-01 | -4.184e-03 | -6.124e-04 |
| 63 | rs419925 | 0.29 | 0.78 | 0.02 | -31953 | 5.128e-01 | 9.761e-01 | -4.210e-03 | -2.839e-04 |
| 64 | rs2075649 | 0.42 | 0.96 | 0.04 | -16749 | 5.640e-01 | 9.829e-01 | -3.684e-03 | -2.228e-04 |
| 65 | rs370705 | 0.29 | 0.78 | 0.02 | -32441 | 4.952e-01 | 9.952e-01 | -4.385e-03 | 5.711e-05 |
| 61 | rs369599 | 0.29 | 0.78 | 0.02 | -32743 | 0.56 | 0.92 | -0.00 | -0.00 |
| 62 | rs3837923 | 0.29 | 0.78 | 0.02 | -32513 | 0.52 | 0.95 | -0.00 | -0.00 |

Continued on next page

**Table S2 continued from previous page**

| | SNP | MAF | $D'$ | $R^2$ | Distance | p: Indirect | p: Interaction | $\hat{\beta}_D$ | $\hat{\beta}_{GE}$ |
|---|---|---|---|---|---|---|---|---|---|
| 63 | rs419925 | 0.29 | 0.78 | 0.02 | -31953 | 0.51 | 0.98 | -0.00 | -0.00 |
| 64 | rs2075649 | 0.42 | 0.96 | 0.04 | -16749 | 0.56 | 0.98 | -0.00 | -0.00 |
| 65 | rs370705 | 0.29 | 0.78 | 0.02 | -32441 | 0.50 | 1.00 | -0.00 | 0.00 |

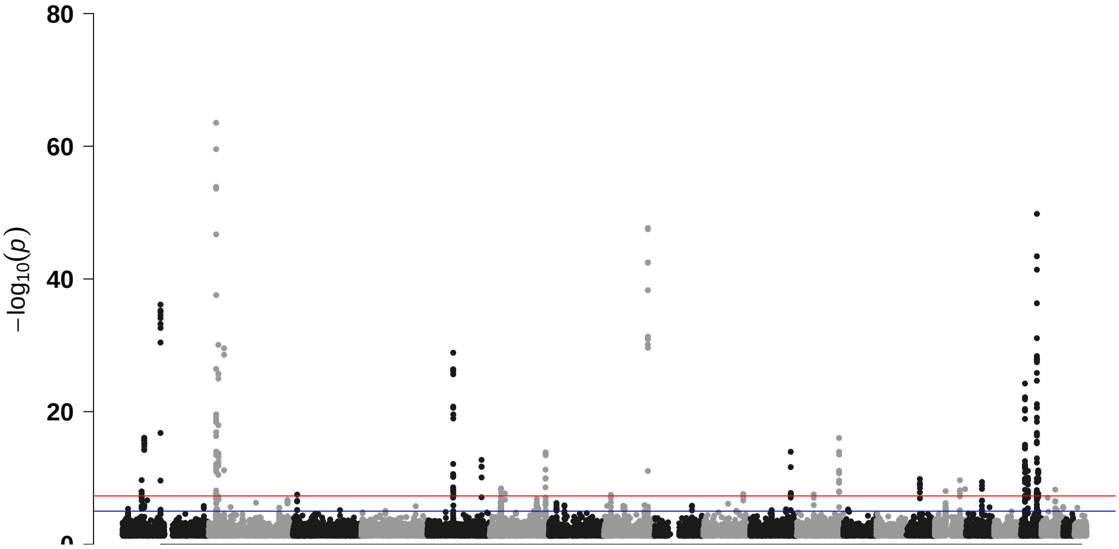


Figure S8: GWAS result using the traditional additive test for the UKB application. The red line denotes the genome-wide significance level of 5e-8.
